# Sexual Dimorphism, Altered Hippocampal Glutamatergic Neurotransmission and Cognitive Impairment in APP Knock-In Mice

**DOI:** 10.1101/2023.12.05.570100

**Authors:** Caleigh A. Findley, Samuel .A. McFadden, Tiarra Hill, Mackenzie R. Peck, Kathleen Quinn, Kevin N. Hascup, Erin R. Hascup

## Abstract

**Background:** It is well established that glutamatergic neurotransmission plays an essential role in learning and memory. Previous studies indicate that glutamate dynamics shift with Alzheimer’s disease (AD) progression, contributing to negative cognitive outcomes.

**Objective:** In this study, we characterized hippocampal glutamatergic signaling with age and disease progression in a knock-in mouse model of AD (APP^NL-F/NL-F^).

**Methods:** At 2-4 and 18+ months old, male and female APP^NL/NL^, APP^NL-F/NL-F^, and C57BL/6 mice underwent cognitive assessment using Morris water maze (MWM) and Novel Object Recognition (NOR). Then, basal and 70 mM KCl stimulus-evoked glutamate release was measured in the dentate gyrus (DG), CA3, and CA1 regions of the hippocampus using a glutamate-selective microelectrode in anesthetized mice.

**Results:** Glutamate recordings support elevated stimulus-evoked glutamate release in the DG and CA3 of young APP^NL-F/NL-F^ male mice that declined with age compared to age-matched control mice. Young female APP^NL-F/NL-F^ mice exhibited increased glutamate clearance in the CA1 that slowed with age compared to age-matched control mice. Male and female APP^NL-F/NL-F^ mice exhibited decreased CA1 basal glutamate levels, while males also showed depletion in the CA3. Cognitive assessment demonstrated impaired spatial cognition in aged male and female APP^NL-F/NL-F^ mice, but only aged females displayed recognition memory deficits compared to age-matched control mice.

**Conclusions:** These findings confirm a sex-dependent hyper-to-hypoactivation glutamatergic paradigm in APP^NL-F/NL-F^ mice. Further, data illustrate a sexually dimorphic biological aging process resulting in a more severe cognitive phenotype for female APP^NL-F/NL-F^ mice than their male counterparts. Research outcomes mirror that of human AD pathology and provide further evidence of divergent AD pathogenesis between sexes.

## Introduction

Glutamate is the most abundant neurotransmitter in the brain, expressed by the vast majority of excitatory neurons ^1^. It is a central component of many neurological functions, including learning and memory ^2^. Glutamate dynamics are impacted by aging-related cortical thinning and synaptic pruning that occurs in the hippocampus and prefrontal cortex, amongst other areas ^3,4^. In the human condition, aging diminishes glutamate levels over time, impacting pivotal processes for learning and memory and contributing to cognitive decline commonly associated with age ^3–5^.

Many pathological conditions involve dysregulation of the glutamatergic system, including epilepsy and aging-related neurodegenerative disorders like Alzheimer’s disease (AD). These conditions include loss of glutamatergic homeostasis that results in neuronal death and brain atrophy ^6^. Glutamatergic hyperactivity has been characterized in both AD mouse models and human pathology, first appearing before the onset of plaque deposition and cognitive symptomology. Prior work from our laboratory demonstrated a hippocampal glutamatergic imbalance that appears in young (2-6 months old) male AβPP/PS1 transgenic AD mice, and shifts in a temporal-specific manner with disease progression ^7^. The CA1 and dentate gyrus (DG) subregions displayed glutamatergic hyperactivity that continued into later disease stages, with a significant drop-off around 18 months of age in the CA1. The pattern found in the CA1 reflects the hyper-to-hypoactivation hypothesis, describing early overstimulation of glutamatergic receptors and astrocytic glutamate transporters leading to dyshomeostasis at the glutamatergic synapse in AD pathology ^8–10^. As well, our laboratory found increased basal glutamate levels in the CA1, CA3, and DG of AβPP/PS1 male mice starting at 6 months of age and continued throughout disease progression. These findings provided further evidence supporting the loss of hippocampal signal-to-noise ratio with disease progression in AD pathology. Signal-to-noise ratio describes the threshold by which neuronal stimulation must overcome to evoke an action potential. An imbalance in this ratio indicates dyshomeostasis at the glutamatergic synapse and can hinder long-term potentiation ^8,9^. This study aims to characterize hippocampal glutamate dynamics in a novel knock-in mouse model of AD (APP^NL-F/NL-F^). The APP^NL-F/NL-F^ mouse model may better recapitulate the human condition and avoids confounding variables associated with transgenic mice, such as overexpression of the amyloid precursor protein and presenilin-1 ^11^. APP^NL-F/NL-F^ mice have a knock-in APP gene with an endogenous promoter and humanized amyloid-beta (Aβ) sequence. The Aβ sequence used also appears in most human sporadic and familial AD patients ^11^. There is no genetic overexpression, however, this model does have the Swedish mutation to APP (NL) that raises total Aβ levels. APP has the Iberian mutation (F), which increases the Aβ42: Aβ40 ratio. This model typically shows plaque deposition at around 6 months old, with steady plaque accumulation up to 18 months old ^12–14^.

To our knowledge, glutamatergic neurotransmission throughout disease progression has not previously been characterized in this model. Further, investigation of sex differences for AD mice is limited ^12,14,15^. Amyloidosis throughout disease progression has been observed as similar between male and female APP^NL-F/NL-F^ mice, while reports of cognitive symptomology yield mixed results ^14,15^. Prior work from our laboratory investigating glutamate dynamics has been limited to male transgenic (AβPP/PS1) mice ^7^. Thus, it remains unclear how sex may impact glutamate and cognition with age and disease progression in this model.

To address this question, we utilized glutamate-specific *in vivo* microelectrode array (MEA) recordings in the hippocampus of APP^NL-F/NL-F^ mice at young (2-4 months old) and aged (18+ months old) time points. The APP^NL/NL^ control model that expresses only the APP Swedish mutation was also examined. Comparing results from the APP^NL-F/NL-F^ model to the APP^NL/NL^ model allows for the elimination of confounding variables such as non-Aβ fragments of APP, including C-terminal fragments ^11^. We hypothesized that APP^NL-F/NL-F^ mice would exhibit presymptomatic hippocampal glutamatergic hyperactivation that shifts with age and disease progression.

## Materials and Methods

### Animals

Founder male and female C57BL/6 (RRID:IMSR JAX:000,664) were obtained from Jackson Laboratory (Bar Harbor, ME). Founder APP^NL/NL^ (RRID:IMSR_RBRC06342) and APP^NL-F/NL-F^ (RRID: IMSR_RBRC06343) male and female mice were obtained from Riken (Tokyo, Japan). Founder mice of all strains were used to start breeding colonies of the respective genotypes and their progeny were used for all experiments. Protocols for animal use were approved by the *Institutional Animal Care and Use Committee* at Southern Illinois University School of Medicine, which is accredited by the Association for Assessment and Accreditation of Laboratory Animal Care. The study was not pre-registered. Mice were group housed according to sex and genotype on a 12:12 h light-dark cycle, and food and water were available *ad libitum*. All experiments were conducted during the light phase. Mouse cages were pseudo-randomized using the Microsoft Excel 2013 randomization function to generate random decimal numbers between 0 and 1 for each mouse cage. No exclusion criteria were pre-determined. 164 animals were used in this study including the following groups: 2-4 months old males (C57BL/6 (n=13), APP^NL/NL^ (n=14), APP^NL-F/NL-F^ (n=16)) and females (C57BL/6 (n=14), APP^NL/NL^ (n=13), APP^NL-F/NL-F^ (n=12)); 18+ months old males (C57BL/6 (n=14), APP^NL/NL^ (n=12), APP^NL-F/NL-F^ (n=15)) and females (C57BL/6 (n=15), APP^NL/NL^ (n=13), APP^NL-F/NL-F^ (n=13). All mice underwent cognitive assessment and *in vivo* glutamate recordings with the exception of electrode failure or death before or during glutamate recordings. Immediately following anesthetized glutamate recordings, mice were euthanized with an overdose of isoflurane followed by rapid decapitation. Genotypes were confirmed by collecting a 5 mm tail tip for analysis by TransnetYX, Inc (Cordova, TN). All mice were tattooed with unique numerical identifiers to blind researchers throughout experimental paradigms.

### 8-Day Hidden Platform Morris Water Maze (MWM)

The MWM examines spatial learning and long-term memory through requiring mice to utilize external visual cues to locate the hidden platform (submerged 1 cm below the opaque water surface), regardless of starting position in the pool. The MWM consists of one acclimation day, five consecutive learning days, and a delayed probe challenge. The five learning days include three trials, up to 90-seconds in duration starting from three different entry points into the pool (order is randomized throughout the learning days) with at least a 20-minute inter-trial-interval to test spatial learning capabilities. After two days without testing mice then undergo a probe challenge consisting of one 60-second trial with the platform removed from the pool to test long-term spatial memory recall. The ANY-maze video tracking system (Stoelting, Wood Dale, IL) tracks and analyzes several parameters throughout the learning days and probe challenge, including average speed, cumulative distance from platform, platform entries, latency to first platform entry, and path efficiency to first platform entry.

### Novel Object Recognition (NOR)

NOR is designed to investigate object recognition memory by measuring the exploration time of a novel object compared to a familiar object within a testing arena. One week after the MWM probe challenge, mice were habituated to the NOR testing arena for 30 minutes. Familiarization training occurred 24 hours later in which two similar objects are placed into the arena and the mice were allowed to explore for 5 minutes. After a 24-hour inter-session-interval, mice were placed back into the arena for 5 minutes with one familiar object from the previous training day and a novel object to test retention. The ANY-maze video tracking system was utilized to track animals throughout the three experiment days and provides measures of distance traveled, average speed, and exploration time to determine the novel object discrimination index.

### *In Vivo* Electrochemistry

Enzyme-based MEAs with platinum (Pt) recording surfaces were prepared for *in vivo* glutamate measurements and calibrated through the FAST-16 recording software as previously described ^7,16,17^. A glutamate oxidase (GluOX) solution was applied to a Pt recording surface to allow for enzymatic degradation of glutamate to α-ketoglutarate and the electroactive reporter molecule H2O2. The adjacent self-referencing recording surface was only coated with an inactive protein matrix and cannot enzymatically generate H2O2 from l-glutamate. The FAST-16 system utilizes the self-referencing site for offline subtraction from the GluOX coated site as a localized control mechanism. A potential of + 0.7 V vs an Ag/AgCl reference electrode was applied to the recording sites for oxidation of H2O2. Recording sites were then electroplated with 5 mM 1,3 phenylenediamine dihydrochloride (mPD) in 0.05M phosphate buffered saline 72 hours after enzymatic coating. mPD provides an exclusion layer that blocks the detection of possible interferants that are electrochemically active at + 0.7 V. MEA calibration occurred on experiment day, with an average ± standard error of the mean (SEM) glutamate sensitivity of 4.8610 ± 0.0002 pA/μM (R^2^ = 0.998 ± 0.001), selectivity ratio of 126 ± 11 to 1, and limit of detection of 0.9305 ± 0.0652 μM based on a signal-to-noise ratio of 3. A micropipette (inner diameter ∼20µm) was then waxed on to the MEA (50-100µm from the electrode surface) for local application of 70mM KCl. Mice were placed into a stereotaxic frame under isoflurane anesthesia and a craniotomy was performed for insertion of the MEA assembly into the hippocampus. Body temperature was maintained with a heated recirculating bath pump attached to a water pad. The starting hemisphere and hippocampal subregion was randomized across the CA1 (AP: −2.0, ML: ±1.0, DV: −1.7 mm) or CA3 (AP: −2.0, ML: ±2.0, DV: −2.2 mm)]. After CA1 recordings, the MEA was lowered into the DG (AP: −2.0, ML: ±1.0, DV: −2.2 mm). MEAs were allowed to baseline for 60 minutes after initial insertion before a 10 s basal glutamate determination and pressure ejection studies commenced. An additional 20-minute baseline period occurred between each hippocampal subregion. Pressure ejections involved a constant volume (∼100-200nL) of 70mM KCl locally applied using a Picospritzer (Parker Hannafin, Morton, IL) attached to the micropipette to evoke glutamate release.

### Cresyl Violet Staining

Mice were euthanized after stimulation by isoflurane overdose and rapid decapitation with sharp scissors. The brain was extracted and fixed in 4% paraformaldehyde for 48 hours and then transferred to 30% sucrose for storage. A cryostat (Model HM525 NX, Thermo Fisher Scientific) was used to obtain 20-micron coronal sections throughout the hippocampus. Slices were mounted on a glass slide, stained with cresyl violet, and coverslipped to verify MEA placement for each mouse.

### Statistical Analysis

Sample size was determined based on previous MWM and electrochemical data using multiple mouse models. GraphPad Prism 9 Software (La Jolla, CA; RRID:SCR 002798) was used for statistical analyses. Statistical tests are listed in each figure legend. Data were not assessed for normality. A single Grubb’s test (alpha = 0.05) was utilized to identify significant outliers in each group. Data are represented as mean ± SEM and statistical significance was defined as p< 0.05.

## Results

### Alterations to glutamatergic neurotransmission before cognitive symptoms in young AD mice

At 2-4 months of age, APP^NL/NL^, APP^NL-F/NL-F^, and C57BL/6 mice underwent a cognitive battery to assess spatial navigation and retention memory. As expected, no significant genotype differences were observed for either sex in spatial learning and memory (Supplementary Figure 1A-E). During the NOR retention day, male mice showed no genotype differences in exploratory behavior and retention memory (Supplementary Figure 1G-I). A main effect for genotype was observed for distance travelled (*F*(2.36)= 12.90, p<0.0001) and average speed (*F*(2.36)= 12.72, p<0.0001) in female mice. Female APP^NL-F/NL-F^ mice showed significantly increased distance travelled and average speed compared to age-matched APP^NL/NL^ (p<0.0001; p=<0.0001) and C57BL/6 mice (p=0.0066; p=0.0068), respectively, that was only exhibited during the retention day. No genotype differences were observed in retention memory for female mice. These findings indicate elevated exploratory behavior in female APP^NL-F/NL-F^ mice during cognitive testing in agreement with previous studies ^15^.

To characterize possible changes in hippocampal glutamate dynamics before cognitive decline, *in vivo* glutamate recordings were conducted on anesthetized mice following completion of the behavioral assays. For male mice, representative traces are shown in Figure 1A. No significant differences were observed in basal glutamate levels for any hippocampal subregion (Figure 1B). However, stimulation of surrounding neurons with 70mM KCl yielded a significant main effect for evoked glutamate release in the DG (*F*(2,29)= 11.01, p=0.003) and CA3 (*F*(2,26)= 6.920, p=0.0039) for male mice (Figure 1C). APP^NL-F/NL-F^ male mice showed elevated stimulus-evoked glutamate release compared to age-matched APP^NL/NL^ (p=0.0016, p=0.0128) and C57BL/6 mice (p=0.0008, p= 0.0087) in the DG and CA3. Time to 80% signal decay (T80) was used to characterize glutamate clearance after stimulation, an important factor for maintaining glutamatergic homeostasis ^18–20^. Examination of T80 glutamate clearance indicated no significant differences for male mice in any hippocampal subregion. These findings demonstrate early hyperactivation of glutamatergic neurotransmission prior to the onset of cognitive symptoms in male APP^NL-F/NL-F^ mice.

**Figure 1:**
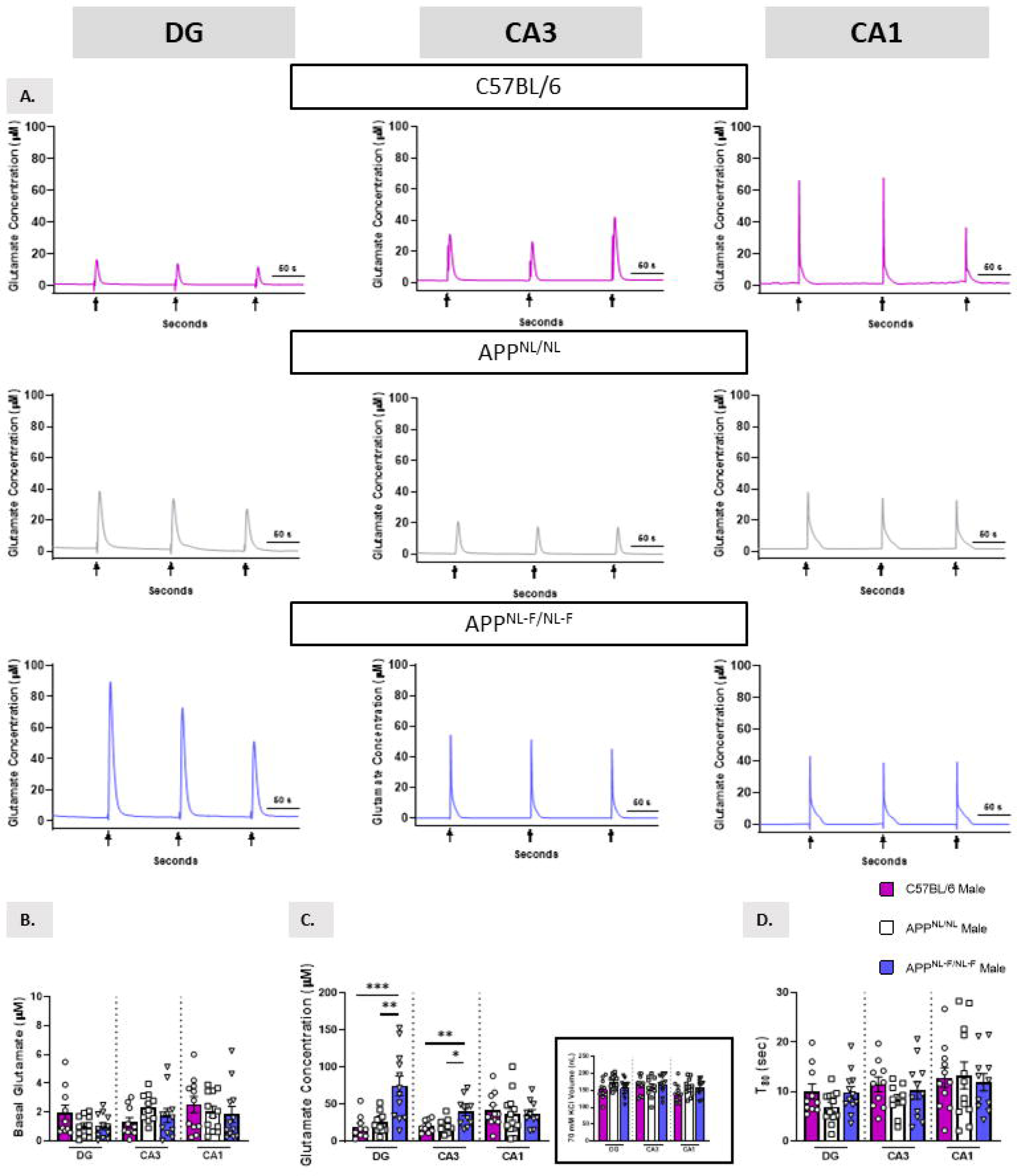
Elevated evoked glutamate release in 2-4 months old APP^NL-F/NL-F^ male mice. A) Representative traces of 70 mM KCl stimulus-evoked glutamate recordings in the DG, CA3, and CA1 hippocampal subregions. B-C) Basal and stimulus-evoked glutamate concentration by subregion. Inset depicts average amount of 70 mM KCl locally applied to evoke glutamate release. D) Time to 80% of glutamate clearance. One-way ANOVA, Tukey’s post-hoc, *p<0.05, **p<0.01.

Representative traces for young female mice are shown in Figure 2A. No significant differences in basal glutamate levels or stimulus-evoked glutamate release for any hippocampal subregion were observed (Figure 2B-C). Analysis of glutamate signal decay time yielded a significant main effect in the CA1 alone for female mice (*F*(2,28)= 4.577, p=0.0191) (Figure 2D). APP^NL-F/NL-F^ females exhibited faster T80 clearance of glutamate compared to age-matched APP^NL/NL^ (p=0.0324) and C57BL/6 mice (p=0.0390). Data support potentially altered glutamate clearance in the CA1 of young female APP^NL-F/NL-F^ mice that may act as an early compensatory mechanism to prevent hyperactivity.

**Figure 2:**
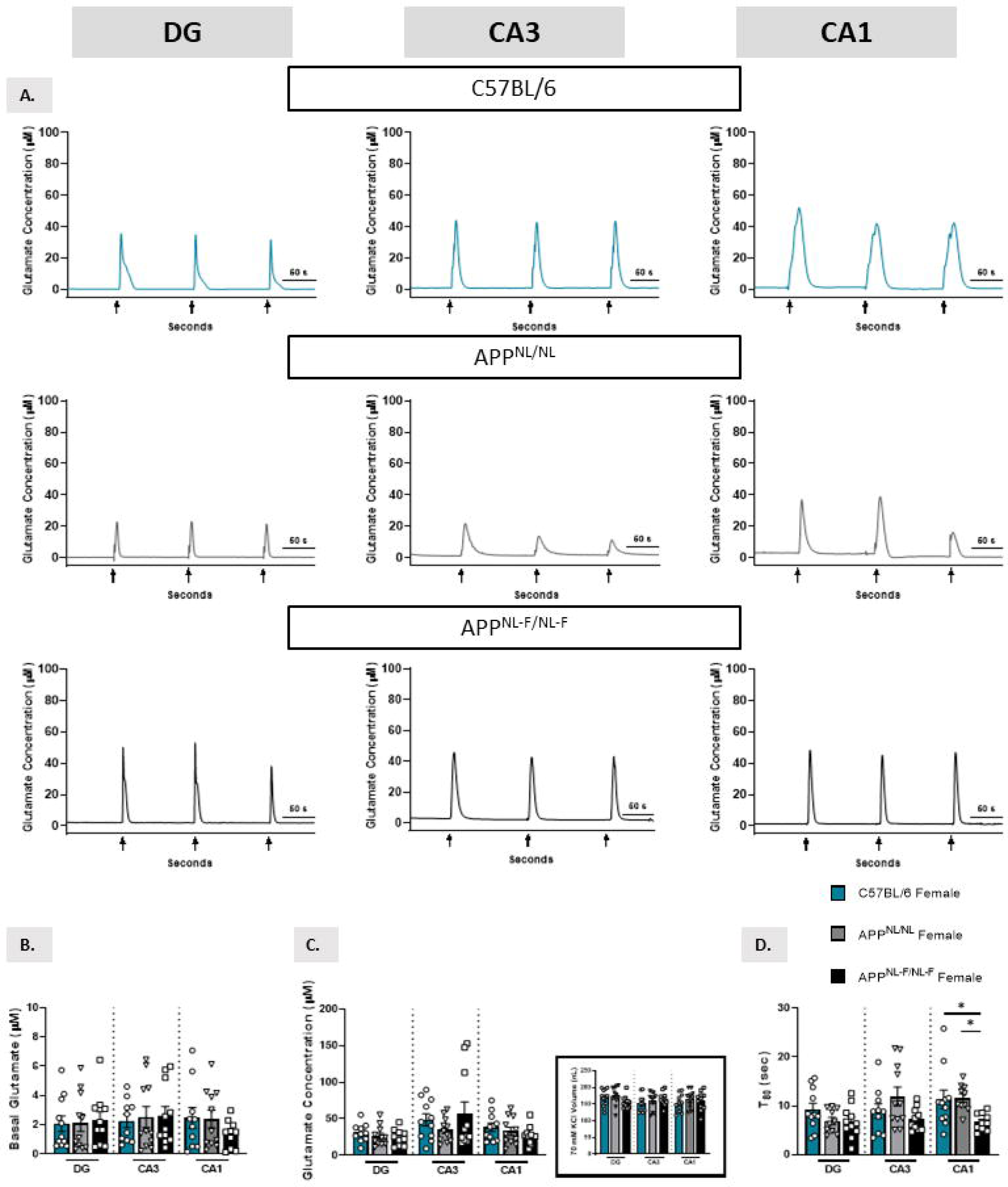
Altered glutamate clearance in the CA1 of 2-4 months old female APP^NL-F/NL-F^ mice. A) Representative traces of 70 mM KCl stimulus-evoked glutamate recordings in the DG, CA3, and CA1 hippocampal subregions. B-C) Basal and stimulus-evoked glutamate concentration by subregion. Inset depicts average amount of 70 mM KCl locally applied to evoke glutamate release. D). Time to 80% of glutamate clearance. One-way ANOVA, Tukey’s post-hoc, *p<0.05.

### Aged AD mice exhibit sex-dependent selective cognitive decline

Previous literature indicates progressive cognitive decline in APP^NL-F/NL-F^ mice, with modest deficits in spatial reversal learning and place preference learning observed at 13-17 months ^12^ and spatial working memory at 18 months ^11^. At 18+ months old, no significant differences were observed for male mice during the learning trials (Figure 3A). A main effect for genotype was observed with cumulative distance from the platform (*F*(2, 38)= 9.828, p= 0.0004) and area under the curve (AUC) (*F*(2,38)= 9.322, p=0.0005) in female mice (Figure 3B). AUC for the five trial days was significantly elevated for female APP^NL-F/NL-F^ mice compared to age-matched APP^NL/NL^ (p=0.0331) and C57BL/6 mice (p=0.0003). Overall, female APP^NL-F/NL-F^ mice had a slower spatial learning curve compared to age-matched C57BL/6 mice. By the final training session, all mice had learned the location of the hidden escape platform. During the probe challenge that is used to evaluate long-term memory recall, a significant main effect was observed for cumulative distance from the platform in female mice (*F*(2,38)= 7.716, p=0.0015). Female APP^NL-F/NL-F^ mice exhibited increased cumulative distance compared to age-matched APP^NL/NL^ (p= 0.0064) and C57BL/6 mice (p=0.0029) (Figure 3C). No significant differences in cumulative distance were observed for male mice. However, analysis of platform entries per distance swam indicated a significant main effect for male mice (*F*(2,38)= 5.978, p=0.0055). APP^NL-F/NL-F^ (p=0.0043) male mice showed a significant decrease in number of platform entries compared to age-matched C57BL/6 mice (Figure 3D). No significant genotype differences were observed for female mice on platform entries. Path efficiency to first platform entry was also examined and yielded no significant findings for either sex (Figure 3E). Representative path traces are shown in Figure 3F. These findings indicate sex differences in spatial learning and memory impairment with age in APP^NL-F/NL-F^ mice, in agreement with prior aging literature examining this mouse model ^14^.

**Figure 3:**
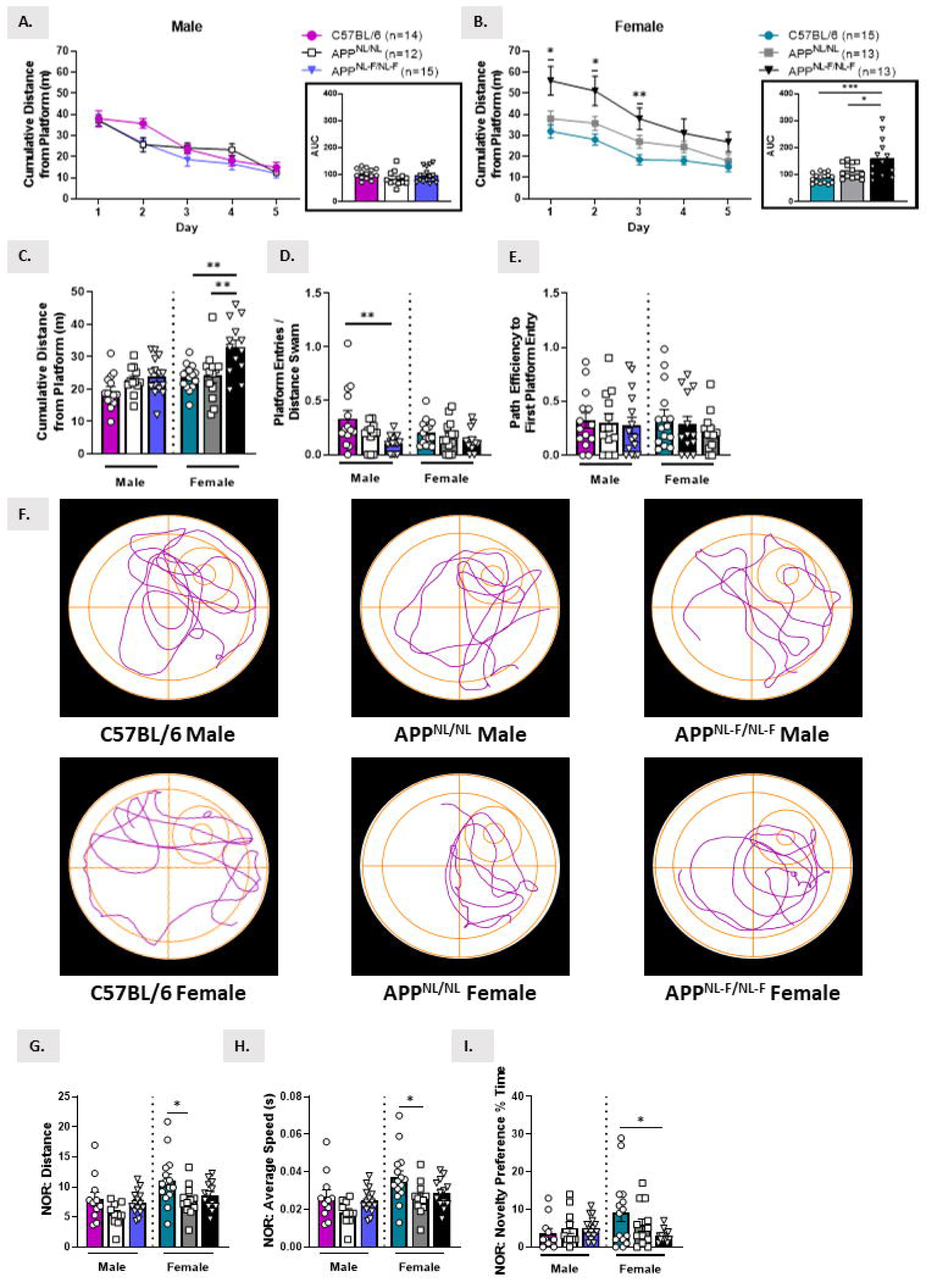
Sex-dependent cognitive decline in aged AD mice. A-B) Cumulative distance to the platform throughout the five trial days for male (A) and female (B) mice. Insets depict area under the curve (AUC) for the five-day learning period. C-E) Probe challenge measures include: cumulative distance, platform entries per distance swam, and path efficiency to first entry. F) Representative track plots from the probe challenge for each sex and genotype. G-H) Distance and average speed during the novel object retention day. I) Evaluation of retention memory through novelty preference. MWM Trials: Two-way ANOVA, Tukey’s post-hoc; MWM Probe and NOR: One-way ANOVA, Tukey’s post-hoc, *p<0.05, **p<0.01, ***p<0.001.

Data from the NOR task indicated no significant genotype differences on any parameters during the retention day for male mice (Figure 3G-I). Significant main effects were observed for distance (*F*(2,37)= 3.947, p= 0.0279), average speed (*F*(2,37)= 4.006, p= 0.0266), and novelty preference (*F*(2,37)= 3.108, p= 0.0565) in female mice. Female APP^NL/NL^ female mice alone exhibited decreased distance (p= 0.0310) and average speed (p= 0.0266) compared to age-matched C57BL/6 mice (Figure 3G-H). APP^NL-F/NL-F^ female mice showed significantly decreased novelty preference compared to age-matched C57BL/6 mice (p=0.0453) (Figure 3I). Together, data support impaired spatial cognition and selective impairment of recognition memory in aged female APP^NL-F/NL-F^ mice, indicating a potentially more severe cognitive phenotype for aged females than males in the APP^NL-F/NL-F^ mouse model.

### Temporal-specific alterations to hippocampal glutamate dynamics in aged AD mice

*In vivo* glutamate recordings were also conducted in aged mice to investigate possible changes in hippocampal glutamate dynamics with age and disease progression. Representative traces for aged male mice are shown in Figure 4A. Analysis of basal glutamate indicated a significant main effect in the CA3 (*F(*2, 27)= 6.486, p=0.0050) and the CA1 (*F*(2, 25)= 3.437, p=0.0480) for male mice (Figure 4B). APP^NL-F/NL-F^ male mice exhibited decreased basal glutamate levels in the CA3 (p=0.0193) and the CA1 (p= 0.0488) compared to age-matched C57BL/6 male mice. No significant differences in basal glutamate were observed for APP^NL/NL^ mice in any hippocampal subregion. A significant main effect for evoked glutamate release was observed in the DG (*F*(2, 27)= 12.06, p=0.0002) and CA3 (*F*(2, 28)= 5.313, p=0.0111) for male mice (Figure 4C). APP^NL/NL^ (p=0.0006, p=0.0209) and APP^NL-F/NL-F^ (p=0.0009, p=0.0280) male mice showed decreased stimulus-evoked glutamate release in the DG and CA3 compared to age-matched C57BL/6 mice. Analysis of T80 also indicated a significant main effect in the DG for aged male mice (*F*(2, 27)= 3.579, p= 0.0418). APP^NL/NL^ male mice showed increased clearance time compared to C57BL/6 male mice (p= 0.0333; Figure 4D). These findings indicate a potentially subregion-specific hypoglutamatergic environment in the hippocampus of aged male AD mice and effectively demonstrate the hyper-to-hypoactivation phenomenon for glutamatergic neurotransmission throughout disease progression in APP^NL-F/NL-F^ male mice.

**Figure 4:**
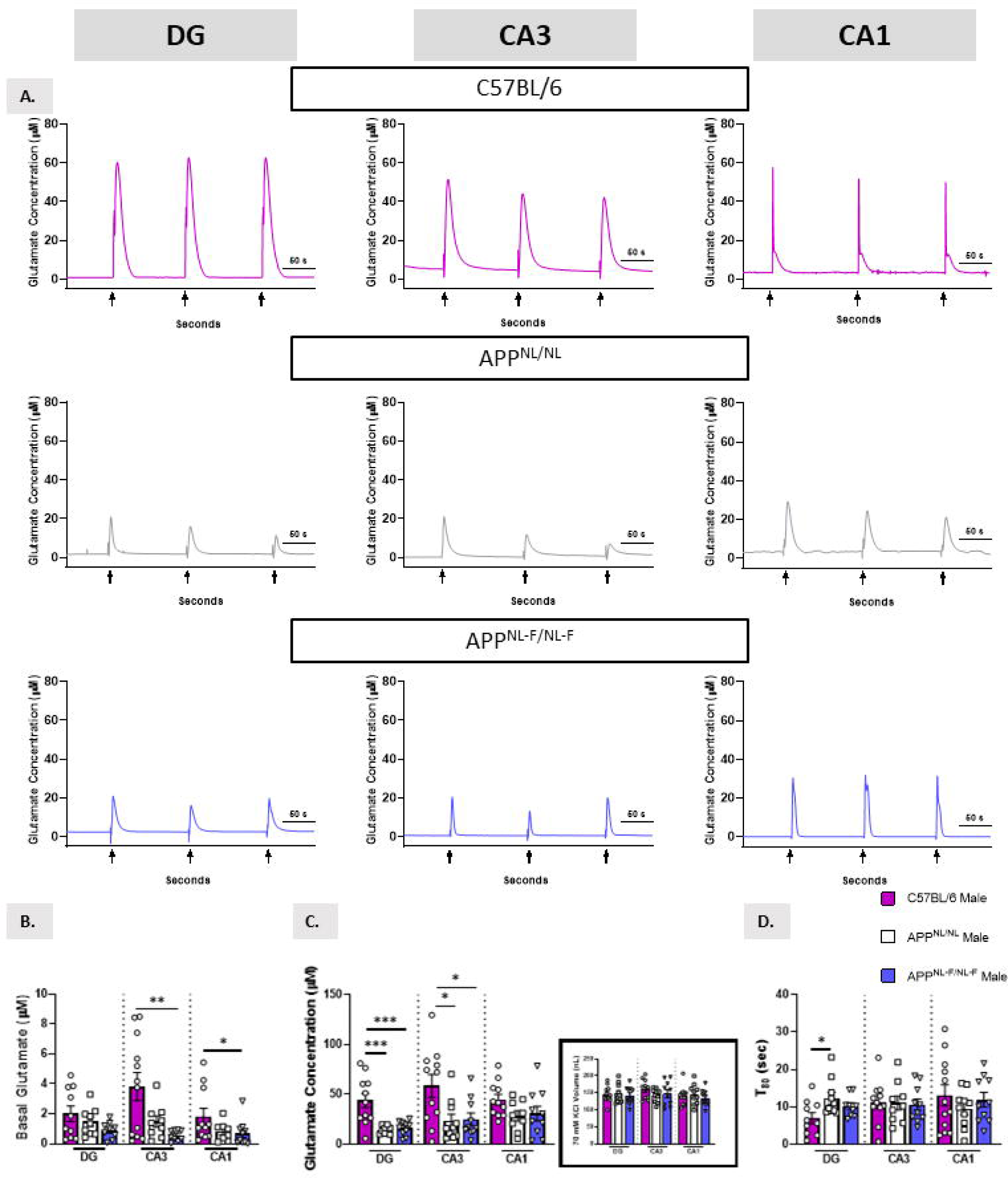
Hypoactive glutamate dynamics in the hippocampus of 18+ months old male AD mice. A) Representative traces from 70 mM KCl stimulus-evoked glutamate recordings in the DG, CA3, and CA1 hippocampal subregions. B-C) Basal and stimulus-evoked glutamate concentration by subregion. Inset depicts average amount of 70 mM KCl locally applied to evoke glutamate release. D) Time to 80% of glutamate clearance. One-way ANOVA, Tukey’s post-hoc, *p<0.05, ***p<0.001.

Representative traces for aged female mice are shown in Figure 5A. A significant main effect (*F*(2, 29)= 5.080, p=0.0128) was observed for basal glutamate levels in the CA1 alone for female mice (Figure 5B). Female APP^NL-F/NL-F^ mice displayed decreased CA1 basal glutamate compared to age-matched C57BL/6 mice (p=0.0093). No differences in basal glutamate were observed for APP^NL/NL^ female mice in any hippocampal subregion. Data support no significant differences in stimulus-evoked glutamate release in any subregion for aged female mice (Figure 5C). A main effect for T80 was observed in the DG for female mice (*F*(2, 27)= 3.579, p=0.0418). APP^NL/NL^ female mice alone exhibited faster T80 than age-matched C57BL/6 mice (p=0.0333) (Figure 5D). Together, data support potential alterations in glutamate clearance and regulation of the glutamatergic homeostasis with age in female APP^NL-F/NL-F^ mice and a unique biological aging process from that observed in male APP^NL-F/NL-F^ mice.

**Figure 5:**
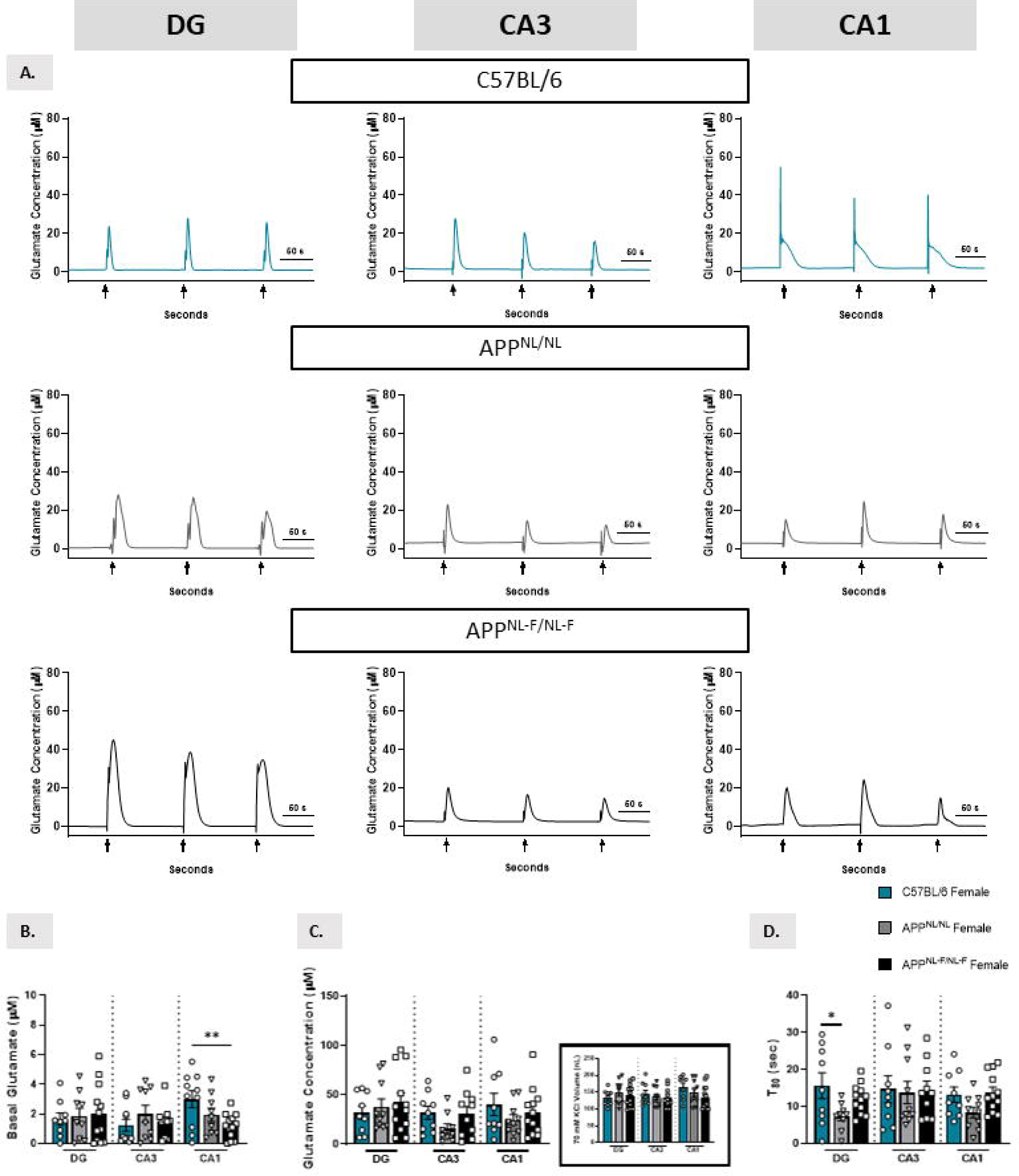
Decreased basal glutamate levels in the CA1 of 18+ months old female APP^NL-F/NL-F^ mice. A) Representative traces from the DG, CA3, and CA1. B-C) Basal and stimulus-evoked glutamate concentration by subregion. Inset depicts average amount of 70 mM KCl locally applied to evoke glutamate release. D) Time to 80% of glutamate clearance. One-way ANOVA, Tukey’s post-hoc, *p<0.05, **p<0.01.

## Discussion

This study aimed to characterize glutamatergic neurotransmission throughout disease progression in the APP^NL-F/NL-F^ mouse model of AD. We utilized a glutamate biosensor to record hippocampal glutamatergic neurotransmission *in vivo* in both sexes at young (2-4 months old) and aged (18+ months old) time points to address this question. Further, we implemented the use of multiple sex- and age-matched C57BL/6 groups, including the APP^NL/NL^ model, to eliminate confounding variables and better analyze the role of Aβ42 pathology. We hypothesized that APP^NL-F/NL-F^ mice would exhibit presymptomatic hippocampal glutamatergic hyperactivation that shifts with age and disease progression. Our findings indicated a more severe cognitive phenotype in aged female APP^NL-F/NL-F^ mice than genotype- and age-matched males, and demonstrated sex differences in hippocampal glutamate dynamics throughout the lifespan. These outcomes reflect that of human AD pathology, in which females bear a heavier disease burden and distinct cognitive profile ^21–23^.

Hippocampal glutamate recordings demonstrated a hyper-to-hypoactivation event for glutamatergic neurotransmission throughout the lifespan of APP^NL-F/NL-F^ male mice. Before the onset of cognitive symptoms, young male APP^NL-F/NL-F^ mice displayed increased evoked glutamate release in the DG and CA3 hippocampal subregions. These were the same subregions in which we observed depleted evoked glutamate release in aged APP^NL-F/NL-F^ male mice. A decrease in basal glutamate in the CA1 and CA3 subregions was also observed in these mice, supporting a hypoactive glutamatergic environment with age and disease progression, similar to what is observed in humans (cite Caleigh’s review manuscript). Loss of glutamatergic stimulation of postsynaptic receptors would impair hippocampal long-term potentiation and confer impaired performance on memory tasks ^7–9^. Soluble Aβ42 appears as a likely driver of hyper-to-hypoactivation, as early hyperactivation was not observed in young male APP^NL/NL^ mice that only express increased total amyloid levels and not specific preference for Aβ42 production. Soluble Aβ42 has been shown to directly interact with the glutamatergic synapse and drive dyshomeostasis with disease progression ^9,24–27^. Further, elevated soluble Aβ42 has been characterized in male APP^NL-F/NL-F^ mice beginning around 2 months of age, concurrent with the window of hyperactivation observed in this study ^11,12^. However, aged male APP^NL/NL^ mice showed decreased evoked glutamate release in the same subregions as age-matched APP^NL-F/NL-F^ mice, leaving open the possibility of multiple drivers for glutamatergic hypoactivation in later disease stages.

Data from glutamate recordings in male APP^NL-F/NL-F^ mice varies from that previously obtained by our laboratory in transgenic AβPP/PS1 male mice ^7^. Namely, we observed a steady incline in basal glutamate levels in all subregions of the hippocampus in AβPP/PS1 male mice that was absent in male APP^NL-F/NL-F^ mice— and juxtaposed for the CA3 and CA1 with age and disease progression. Further, a similar hyper-to-hypoactivation event occurred with disease progression in male AβPP/PS1 mice, but primarily in the CA1 instead of the DG and CA3 as observed in APP^NL-F/NL-F^ male mice. This divergence could be due to differences between the mouse models including gene overexpression and transgenic versus knock-in genetic manipulation. AβPP/PS1 exhibit greater amyloid burden, higher early attrition rate, and more aggressive cognitive decline ^14^. However, previous studies have shown a similar amyloidosis timeline between AβPP/PS1 and APP^NL-F/NL-F^ mice, first appearing in cortex and hippocampus ^7,11,12,14^. Temporal progression through the hippocampus is also similar between the models, occurring first and primarily in the CA1 and then moving into the DG and CA3 ^7,14^. Yet, the temporal progression for amyloidosis and glutamate hyperactivity in male AβPP/PS1 mice mirror each other, unlike that observed in APP^NL-F/NL-F^ mice. Other intermediating factors may come into play – such as mouse model differences in the expression of key glutamate synaptic components across hippocampal subregions – and should be explored in future studies.

Female APP^NL-F/NL-F^ mice underwent a unique and divergent biological aging process to that characterized in male APP^NL-F/NL-F^ mice. No alterations in evoked glutamate release appeared throughout the lifespan for APP^NL-F/NL-F^ female mice — their story pertained more to glutamate clearance and homeostatic maintenance. Young APP^NL-F/NL-F^ female mice showed faster glutamate clearance times compared to age-matched APP^NL/NL^ and C57BL/6 mice in the CA1. The same subregion then showed a loss of basal glutamate levels in aged female APP^NL-F/NL-F^ mice, indicating a possible early compensatory mechanism to maintain glutamatergic homeostasis that failed with disease progression. Glutamatergic transport is the crux of homeostasis at the tripartite glutamate synapse — dysregulation of astrocytic glutamate clearance can significantly impact cognitive outcomes ^28–30^. Of note, a prior study from our laboratory found that local Aβ42 application stimulated lactate release only in the CA1 of female C57BL/6 mice, a possible indication of Aβ42-mediated increased glutamate clearance localized to this hippocampal subregion in female mice ^24^. It is possible that glutamatergic alterations exist more in tonic or spontaneous activity for APP^NL-F/NL-F^ female mice than with stimulus-evoked glutamate release. Such a possibility would explain the absence of effects to evoked glutamate release, and provide more context to faster glutamate clearance times in young APP^NL-F/NL-F^ female mice. Another possible explanation may reside in astrocyte activity and response to pathology, in which prior studies indicate sexual dimorphism and emphasize the neuromodulatory role of estrogen ^31^. Estrogen can protect against glutamate hyperexcitability by supporting astrocyte function and inhibitory regulation ^32,33^ — and yield devastating biological consequences when lost during menopause ^34,35^. However, this is not supported by the results of this study as immunohistochemical evaluation of astrocytes examining both GFAP and GLT-1 did not show significant differences (data not shown). Taken together, there are likely multiple factors contributing to the difference in hippocampal glutamate dynamics with disease progression for female APP^NL-F/NL-F^ mice.

Glutamate findings in female APP^NL-F/NL-F^ mice contrast that of previous studies from other laboratories in female AβPP/PS1 mice ^36^. A loss of hippocampal evoked glutamate response with disease progression was reported, with no alterations to basal glutamate throughout the lifespan. Prior examination of amyloid pathology from both models indicate more aggressive amyloidosis in female AβPP/PS1 mice, with both models following a similar temporal progression observed in their male counterparts ^14^. At 12 months of age, female AβPP/PS1 mice display more severe amyloid pathology, especially in the hippocampus, than male AβPP/PS1 mice ^37^. It is possible that the severity of amyloid pathology observed in female AβPP/PS1 mice drove the glutamatergic hypoactivity observed in later disease stages and would explain the absence of this effect in APP^NL-F/NL-F^ female mice.

Results from our behavioral analysis indicated sexual dimorphism in cognitive deficits for APP^NL-F/NL-F^ aged mice. Aged male APP^NL-F/NL-F^ mice exhibited impaired spatial long-term memory during the MWM, but did not show any loss in recognition memory. These findings are congruent with previous studies examining cognition in APP^NL-F/NL-F^ mice ^11,12,14,38^. Aged APP^NL/NL^ male mice did not exhibit deficits in spatial or retention memory, in agreement with previous studies examining 24-month-old male APP^NL/NL^ mice ^39^. It is possible that prolonged glutamatergic dyshomeostasis in multiple subregions for APP^NL-F/NL-F^ male mice contributed to an earlier onset of cognitive symptoms than APP^NL/NL^ mice. Aged female APP^NL-F/NL-F^ mice showed impaired spatial learning and memory and recognition memory, collectively displaying a more severe cognitive phenotype than that observed in male mice. As previously discussed, this may be due to loss of astrocytic homeostatic regulation of the glutamatergic synapse with disease progression. This finding mirrors previous studies indicating that AD tends to be more aggressive in female AD mice and in humans ^37,40^. Spatial long-term memory impairment for female APP^NL-F/NL-F^ mice was also observed on a different parameter than genotype- and age-matched male mice during the MWM probe challenge. It is possible that sex differences in spatial navigation and wayfinding could have impacted the parameters by which cognitive deficits were observed ^41^. Previous studies would indicate that this divergence between sexes is unlikely to be attributed to sex differences in executive function ^42^. Sexual dimorphism in glutamate and cognition have been previously described ^43–45^, and support that differences in hippocampal glutamate dynamics observed throughout the lifespan may have also influenced sex differences in cognitive decline for this model.

Collectively, these findings indicate genotype and sex differences in glutamate dynamics and cognition for knock-in AD mice. APP^NL/NL^ mice provided a necessary middle ground between C57BL/6 littermates and APP^NL-F/NL-F^ mice to delineate the role of Aβ42 in study outcomes. Further, these genotype differences also support that APP knock-in alone exerts moderate pathology, which is compounded by increased Aβ42 production generating a more severe cognitive profile. Biological differences in sex add on another layer of complexity to the observed findings and point to worsened pathology in female mice likely implicating a hormonal impact to glutamate and cognition with age and disease progression that warrants further study.

Sexually dimorphic rates of biological aging may also contribute to the differences in glutamate dynamics and cognitive performance. Evidence suggests females maintain better cellular health for most of their life, but become frailer when approaching older age than males ^46^. Divergent disease progression between the sexes would hold important implications for therapeutic treatment. Interventional strategies must evaluate opportune windows for treatment that are tailored to the sex-specific degeneration timelines and broader disease pathogenesis to maximize efficacy. Further, these findings also reinforce that experimentation protocols for therapeutic candidates need to include evaluations of male and female participants. It is entirely plausible that intervention strategies may produce variable outcomes between sexes due to hormonal neuromodulation and sex-dependent pathogenic phenotype.

## Abbreviations

(AD): Alzheimer’s disease
(MWM): morris water maze
(NOR): novel object recognition
(DG): dentate gyrus
(Aβ): amyloid-beta
(MEA): microelectrode array
(Pt): platinum
(GluOX): glutamate oxidase
(mPD): 1,3 phenylenediamine dihydrochloride
(SEM): standard error of the mean
(T80): time to 80% signal decay

## Author Contribution

CAF executed electrochemistry recordings, behavioral assays, data analysis and interpretation, and drafting of the manuscript. SAM assisted with conducting behavioral assays.TH assisted with cryosectioning and cresyl violet staining. MRP and KQ managed the breeding colony. KNH and ERH conceived and supervised the study, imparted surgical and electrochemistry training, and provided manuscript revisions.

## Acknowledgements

The authors have no acknowledgments to report.

## Datasets/Data Availability

The data to support the findings of this study are available from the corresponding author upon reasonable request.

---

Supplementary Figure 1: *No memory deficits present in 2-4 months old knock-in AD mice.* A-B) Spatial learning performance evaluated by cumulative distance from the platform area throughout the five trial days for male (A) and female (B) mice. C-E) Long-term spatial memory recall performance during probe challenge was measured by: cumulative distance, platform entries per distance swam, and path efficiency to first entry. G-H) Distance and average speed during the novel object retention day. I) Evaluation of retention memory through novelty preference. MWM Trials: Two-way ANOVA, Tukey’s post-hoc; MWM Probe and NOR: One-way ANOVA, Tukey’s post-hoc, **p<0.01, ****p<0.0001

