## Supplementary figures and images for "Sexual Dimorphism, Altered Hippocampal Glutamatergic Neurotransmission and Cognitive Impairment in APP Knock-In Mice"

### Supplemental Figure 1

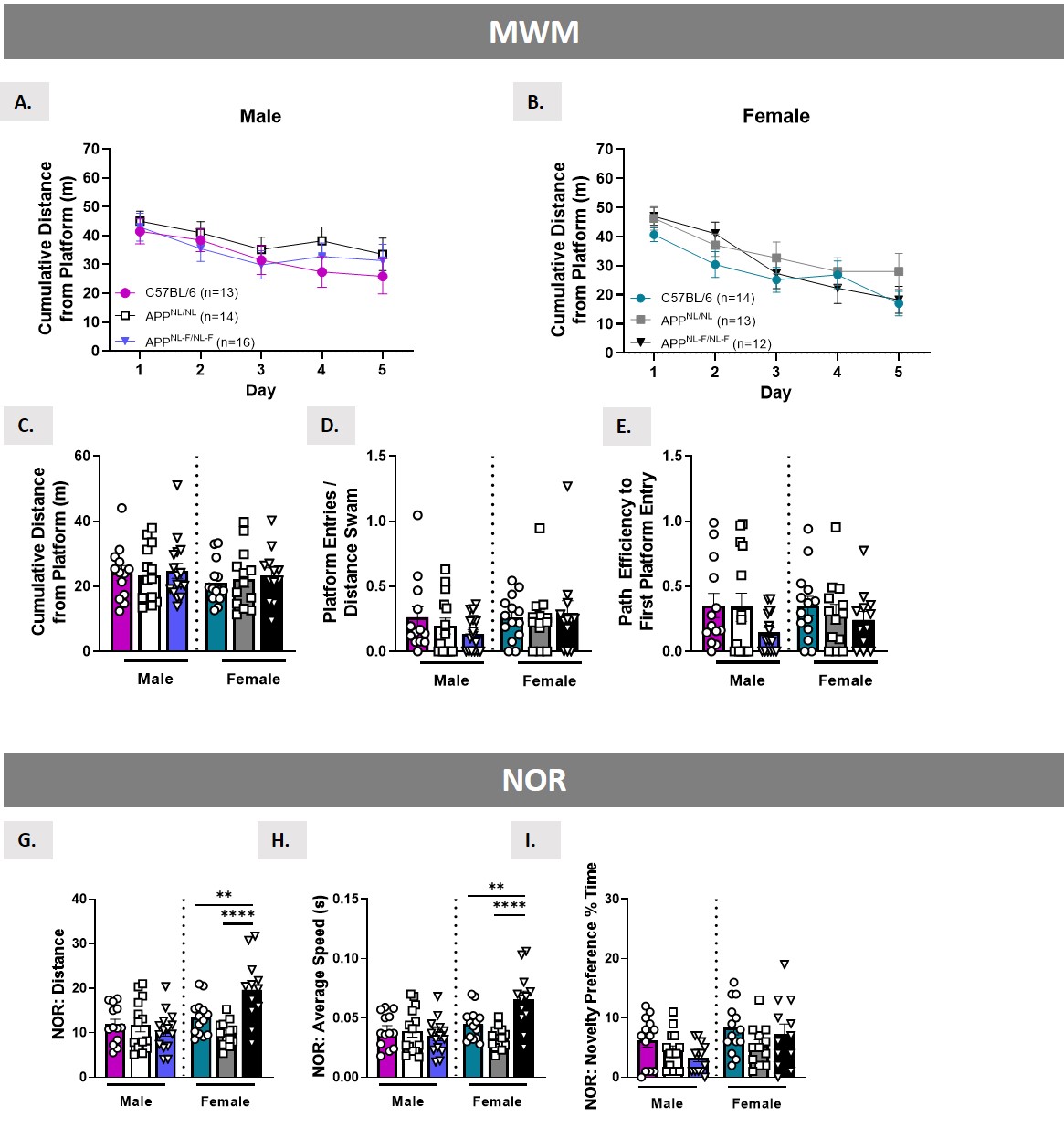
